# MetaX: A peptide centric metaproteomic data analysis platform using Operational Taxa-Functions (OTF)

**DOI:** 10.1101/2024.04.19.590315

**Authors:** Qing Wu, Zhibin Ning, Ailing Zhang, Xu Zhang, Zhongzhi Sun, Daniel Figeys

**Affiliations:** School of Pharmaceutical Sciences, Faculty of Medicine, University of Ottawa, Ottawa, Canada; Department of Biochemistry, Microbiology and Immunology, Faculty of Medicine, University of Ottawa, Ottawa, Canada; Regulatory Research Division, Biologic and Radiopharmaceutical Drugs Directorate, Health Products and Food Branch, Health Canada, Ottawa, Canada

**Author notes:** Correspondence: Daniel Figeys.

**Keywords:** Metaproteomics, MetaX, Peptide-Centric, Operational Taxa-Functions (OTF), Microbial Ecosystems

## Abstract

Metaproteomics analyzes the functional dynamics of microbial communities by identifying peptides and mapping them to the most likely proteins and taxa. The challenge in this field lies in seamlessly integrating taxonomic and functional annotations to accurately represent the contributions of individual microbial taxa to functional diversity. We introduce MetaX, a comprehensive tool for analyzing taxa-function relationships in metaproteomics by mapping peptides to their lowest common ancestors and assigning functions based on proportional thresholds, ensuring accurate peptide-level mappings. Importantly, MetaX introduces the Operational Taxa-Functions (OTF), a new conceptual unit for exploring microbial roles and interactions within ecosystems. Additionally, MetaX extends traditional taxonomic classification by adding a genome level below the species level, enhancing the accuracy of function attribution to specific genomes. We demonstrated MetaX by reanalyzing metaproteomic data from gut microbiomes exposed to various sweeteners, achieving results similar to traditional protein analysis. Furthermore, using the peptide-centric approach and OTF, we observed that *Parabacteroides distasonis* significantly responds to certain sweeteners, highlighting its role in modifying specific metabolic functions. With its intuitive, user-friendly interface, MetaX facilitates detailed study of the complex interactions between microbial taxa and their functions in metaproteomics. It enhances our understanding of microbial roles in ecosystems and health.

## Background

Microbiomes are complex ecosystems important for human health (1–3). Various omics technologies have been introduced to study microbiomes, including 16S rRNA sequencing (4), metagenomics (5), metatranscriptomics (6), and metaproteomics (7). Among these, metaproteomics directly analyzes the presence and abundance of microbial peptides and proteins highlighting the biochemical processes active in the microbiome (8–12). Furthermore, metaproteomics permits simultaneous analysis of both taxonomic distribution and functional status based on the same mass spectrometry dataset.

In metaproteomics, the shotgun approach followed by database searches is commonly used for protein identification and quantification. Unfortunately, in metaproteomics differentiating between highly similar protein sequences becomes challenging as sequence coverage declines. Shared peptides, present in multiple sequence database entries, can sometimes be assigned to a particular protein without a definitive basis, among several potential candidates (14). One prevalent solution is the application of Occam’s razor to generate a minimal list of proteins sufficient for explaining all observed peptides (15). Various tools, like MetaLab (16), employ statistical models to analyze both functions and taxonomies in metaproteomics. Conversely, the anti-Occam’s razor approach aims for a maximal explanatory protein set, in which each protein identified by at least one peptide is included, as utilized in MetaProteomeAnalyzer (17). These approaches have their drawbacks: Occam’s razor may underestimate, while anti-Occam may overestimate the protein diversity in a sample (18).

Another distinct challenge in metaproteomics is the ambiguous association between taxa and functions, often due to peptide sequence conservation across various bacterial taxa. In recent years, peptide-centric methods have been developed to directly annotate functions and taxa from peptides(19–21). MetaGOmics (20) assigns functional and taxonomic annotations to each peptide but limits functional annotation to Gene Ontology (GO). Unipept (19) provides functional data from The UniProt Knowledgebase (UniProtKB) (22) and taxonomic information by calculating the lowest common ancestors (LCA) for peptides. However, it falls short in integrating peptide intensity for cross-sample comparisons and cannot delineate specific functions for each species.

In this study, we introduce MetaX a peptide-based data analysis platform that annotates both taxonomic and functional attributes at the peptide level by employing proportion thresholds. This approach establishes a direct correspondence between function and taxa within the metaproteomic context (“who is doing what and how”). Unlike traditional methods that focus on specific types of functions, our methodology is designed to be universally applicable across all annotatable functional categories. Central to MetaX is the introduction of the Operational Tax-Function (OTF) concept, an operational unit that assigns each peptide a taxa-function between taxonomic identification and functional analysis. MetaX not only streamlines the data analysis process but also facilitates real-time result visualization. The peptide centric approach and the OTF of MetaX allowed novel discovery when reanalyzing metaproteomic datasets previously analyzed by a protein centric approach.

## Methods

The core concept of MetaX is to conserve the information provided by metaproteomics at the peptide level, to annotate the peptides and to provide multiple options for data processing and analysis. Instead of distributing peptides to the most likely proteins, MetaX transfer annotations at the peptide level, allowing comparative analysis of peptide profiles across multiple samples based on their functional annotations, taxonomic annotations, and Operational Taxa-Function. MetaX, written in Python, consists of four key modules: Database Builder, Database Updater, Peptide Annotator, and Operational Taxa-Functions Analyzer. Both command-line interface (CLI) and GUI versions are available for each module.

**Figure 1.**
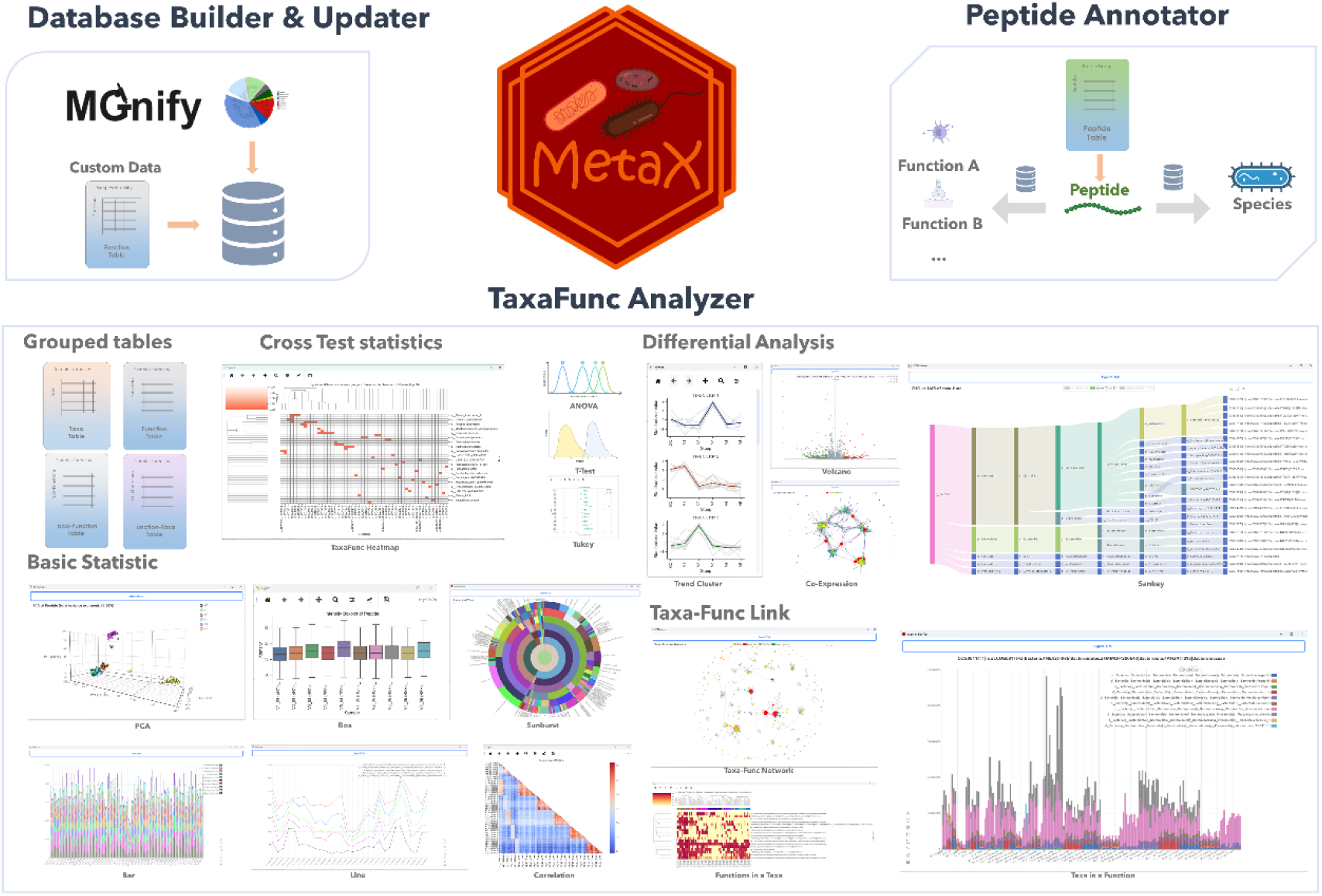
The overview of MetaX.

### Linking Taxa and Functions at the Peptide Level

The association of taxa and functions is realized through peptide-level annotation. Initially, peptides are assigned to a corresponding protein group, incorporating all proteins linked to the peptide. Following this assignment, taxonomic and functional annotations for each peptide are independently conducted.

### Database Construction and Maintenance: Database Builder and Updater

The Database Builder module facilitates the creation of an SQLite database that captures protein function and taxa information. It is preloaded with nine distinct databases from MGnify (23). As well, the Database Builder allows users to formulate databases based on their specific protein data.

Once the foundational SQLite database is established, the Database Updater module provides a functional extension to supplement protein annotations. This module is integrated with several pre-established databases (Human Gut, Human Oral, Cow Rumen and Marine) from dbCAN-seq (24), to streamline the update process. Alternatively, users can implement updates employing their individual data tables.

### Operational Taxa-Functions Table Generation: Peptide Annotator

The Peptide Annotator module incorporates an algorithm to determine the LCA and associated function. This module accepts either a list of proteins or a table delineating the protein groups for each peptide, along with a proportion threshold. Within this module, each protein from a group is queried from the database created by the Database Builder module.

#### Taxonomic Annotation Based on Proportional Threshold

MetaX annotates each peptide based on the annotation information of proteins within its corresponding protein group. It facilitates the determination of the Lowest Common Ancestor (LCA) using a proportional threshold. The taxa proportion for each peptide is then calculated by dividing the occurrence frequency of the most prevalent taxon by the total number of proteins in the group (Fig. 2). This calculation is carried out iteratively, from the genome level up to the domain level, until the taxa proportion meets or exceeds a set threshold. If a peptide does not meet the threshold at the domain level, it is marked at the life level. The default threshold is established at 1, indicating that for a coherent taxonomic annotation, all proteins within a group must be consistently identified with the same taxa at the current level.

**Figure 2.**
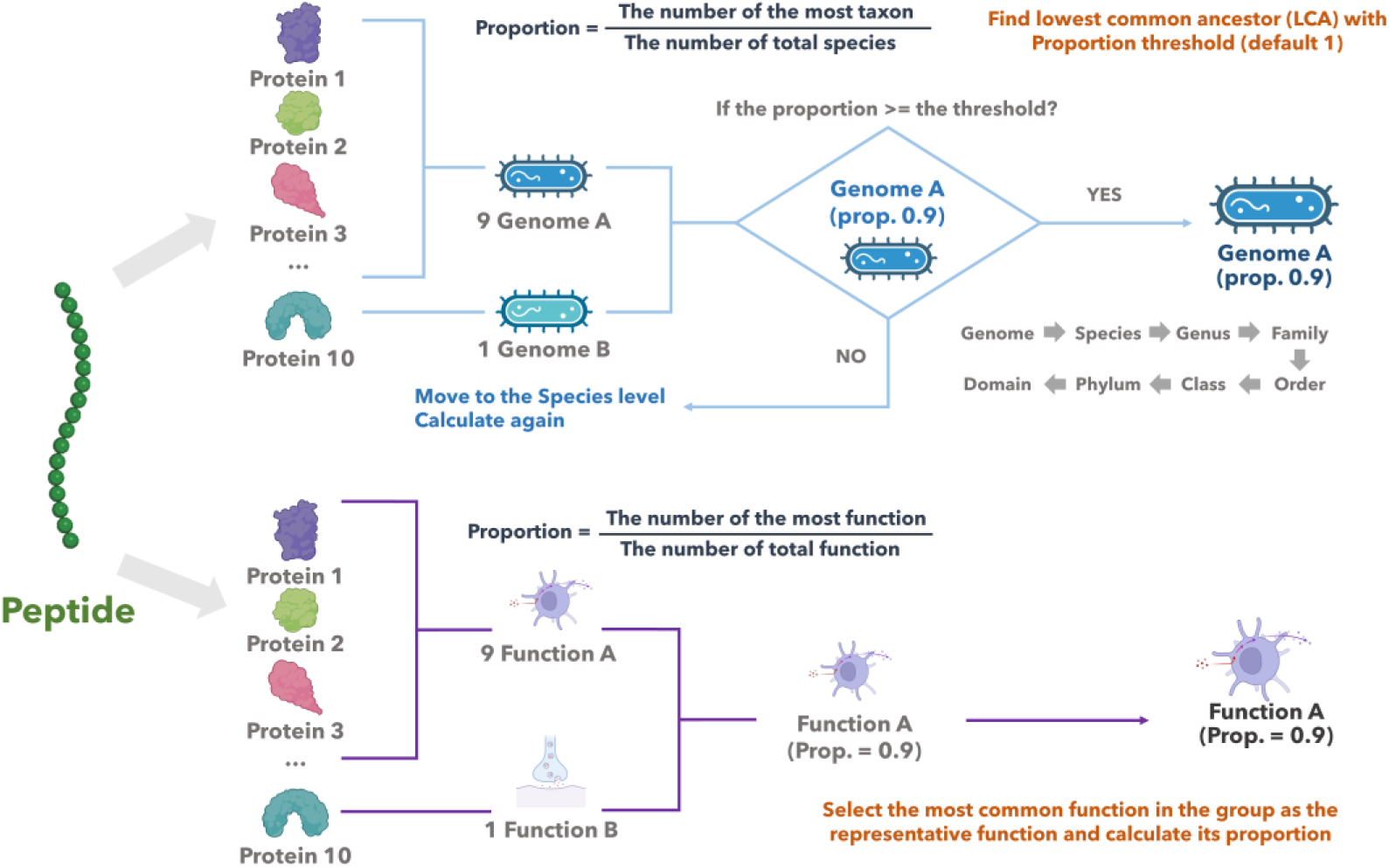
The methodology employed to identify the lowest common ancestors is predicated on the proportional threshold for each peptide, and the methodology for annotating the function of each peptide.

#### Functional Annotation Across Multiple Databases

Functional annotation of each peptide proceeds by identifying the most prevalent function within its associated protein group. Unlike taxonomic annotation, this process does not use a threshold for selection. Instead, the most common function is assigned to the peptide, along with a proportional value indicating its frequency relative to the total number of proteins in the group (Fig. 2). This functional annotation is systematically applied across various databases, such as Gene Ontology (GO) (25), Kyoto Encyclopedia of Genes and Genomes (KEGG) (26), Enzyme Commission number (EC number) (27) and Carbohydrate-Active enZymes (CAZy) databases (28), ensuring a thorough and comprehensive functional profiling of each peptide. This approach facilitates a multi-dimensional understanding of peptide functions, enriching the analysis with diverse functional insights from several established biological databases. For proteins without function annotation, the result returned is “not found”.

### Downstream Analysis: Operational Taxa-Functions Analyzer

The OTF Analyzer module analyses the operational taxa-functions tables, sourced from the Peptide Annotator module, or externally provided by users. Essential table components include peptide information, associated taxa and function annotations, their respective proportions, and peptide intensity. Moreover, users can provide a meta table detailing group information for each sample. If a meta table is missing, MetaX simplifies the process by auto-generating two meta columns, one grouping all samples together and the other assigning each sample to its own category. This approach enables straightforward data analysis, resulting in the creation of an OTF object tailored to diverse analytical needs.

#### Data preprocessing

MetaX offers a series of data preprocessing options including outlier detection, outlier handling, batch effect removal, data transformation, and data normalization. Multiple methods are available for outlier detection, including Interquartile Range (IQR), Half-Zero, Zero-Inflated Poisson, Negative Binomial, Z-Score, and Mahalanobis Distance. These methods are comprehensively detailed in **Table 1 of the Supplementary Document,** and implemented by SciPy (29). Outlier handling can be performed using techniques such as Mean, Median, K-Nearest Neighbors Algorithm (KNN), Regression, and Multiple by using the Python package scikit-learn (30) and Statsmodels (31), as detailed in **Table 2 of the Supplementary Document**. The reCombat (32) the package is leveraged for batch effect removal. Data transformation processes can be done using Log2, Log10, square root transformation, and cube root transformation. NumPy (33) and pandas (34) are used for data normalization, methods include Standard Scaling (Z-Score), Min-Max Scaling, Pareto Scaling, Mean Centering, and Normalization by Sum. Users have the flexibility to select any or all of these methods for data preprocessing, with the additional option to modify the order of data preprocessing steps.

#### Taxonomic Level and Function Setting

MetaX allows users to customize their analyses by choosing specific functions and taxonomic levels. It also allows for the adjustment of function proportion thresholds to suit specific research needs. Upon setting these parameters, MetaX identifies relevant peptides, organizes them into a structured table, and generates four specialized statistical tables: the taxa table, function table, taxa-functions table, and functions-taxa table. Users have the flexibility to establish a minimum count of peptides required for each taxon, function, and taxa-functions, ensuring focused and meaningful analysis. MetaX also provides the option to infer protein intensity tables using either Occam’s razor or the anti-razor method, facilitating detailed downstream analysis.

In the taxa table, peptides associated with the same taxon are consolidated, enabling a more comprehensive understanding of that particular taxon. The function table, on the other hand, merges peptides sharing the same function, which facilitates a more nuanced analysis of that function. The taxa-functions and functions-taxa tables combine different functions within the same taxon or different taxa within the same function, respectively, offering a comparative perspective across taxa and functions.

#### Statistical Analyses and Visualization

Each data table prepared by MetaX is compatible with numerous analytical methods. In particular, Principal Component Analysis (PCA), and technique for variability analysis within large data sets, were performed using the scikit-learn (30) Python library. Alpha and Beta diversity are calculated by scikit-bio (35). The t-test, Dunnett’s test and analysis of variance (ANOVA) tests, which are statistical methods used to compare the means of groups, were conducted using SciPy (29). Tukey’s test, a post hoc analysis performed following an ANOVA test, was carried out with the Statsmodels (31) Python module. The calculation of log2FoldChange with Wald tests between groups, which provides insight into the magnitude and direction of changes, was performed using the PyDESeq2(36) package.

In order to quantify the association between groups, Pearson’s correlation coefficient and Spearman’s rank correlation coefficient were utilized. The former measures linear correlation, whereas the latter assesses monotonic relationships. K-means clustering was employed to identify groups that exhibit similar intensity expression trends, providing further insights into patterns within the data.

All graphical representations of data were created using Seaborn (37), a Python data visualization library based on Matplotlib (38), and Pyecharts (https://github.com/pyecharts/pyecharts/), a Python interactive charting library. These tools provide a wide array of plotting capabilities, allowing for comprehensive graphical exploration of data.

### Graphical User Interface

The GUI of MetaX was developed using PyQt5 (39), a set of Python bindings for Qt libraries that are extensively used to create multi-platform applications. For theming purposes, Qt-Material (https://github.com/UN-GCPDS/qt-material) was employed, enabling the creation of appealing and consistent user interface designs. Furthermore, qtawesome (https://github.com/spyder-ide/qtawesome) was used for the integration of icons in the menu list, enhancing user navigation and the overall aesthetic appeal of the interface.

### Test Dataset

MetaX was tested using raw data files from 175 samples sourced from a previous research study by Sun et al. (40). These raw data files were processed through the MetaLab-MAG (16) software suite with Unified Human Gastrointestinal Protein (UHGP) database V2.1 from MGnify.(23) Ultimately, the resultant ‘final_peptide.tsv’ table, generated via this process, was employed as the testing dataset for evaluating the performance of MetaX.

## Results

### Enhancing Metaproteomic Analysis with Operational Taxa-Functions

MetaX is a peptide-based data analysis platform that introduces the notion of Operational Taxa-Functions (OTF), a conceptual framework that groups peptides by their shared taxonomic and functional attributes. This approach organizes peptides based on their taxonomic identities and functional capacities facilitating the study of the interactions between microbial taxa and their functions. Additionally, MetaX was developed with the goal of maximizing the customizability based on user needs by providing access to multiple tools for processing and analyzing data. To accommodate researchers who are not familiar with command-line operations, the GUI version of MetaX integrates all data processing and analysis.

MetaX is equipped with a broad array of statistical and visualization tools designed to facilitate the in-depth analysis of biological data, including PCA, 3D PCA, Correlation Plots, Box Plots, Heat Maps, Bar Plots, and Line Plots. These tools serve the analysis of peptides, taxa, functions, OTFs, and proteins, enabling researchers to visually explore the data’s complexity. Additionally, MetaX offers specialized tools for taxa analysis such as Alpha-diversity, Beta-diversity, and Tree Maps, along with Sankey Plots for investigating taxa and function relationships. The platform’s filter function simplifies the identification of key items for analysis based on various metrics, such as intensity or frequency across samples. Further customization is facilitated through adjustable parameters for visualization, including size, color, scale, and clustering, providing a versatile environment for detailed analytical exploration (Fig. 3a). MetaX also supports an array of statistical tests, including T-tests, ANOVA, Dunnett’s test, and Tukey’s test, for comprehensive analysis across different datasets.

**Figure 3.**
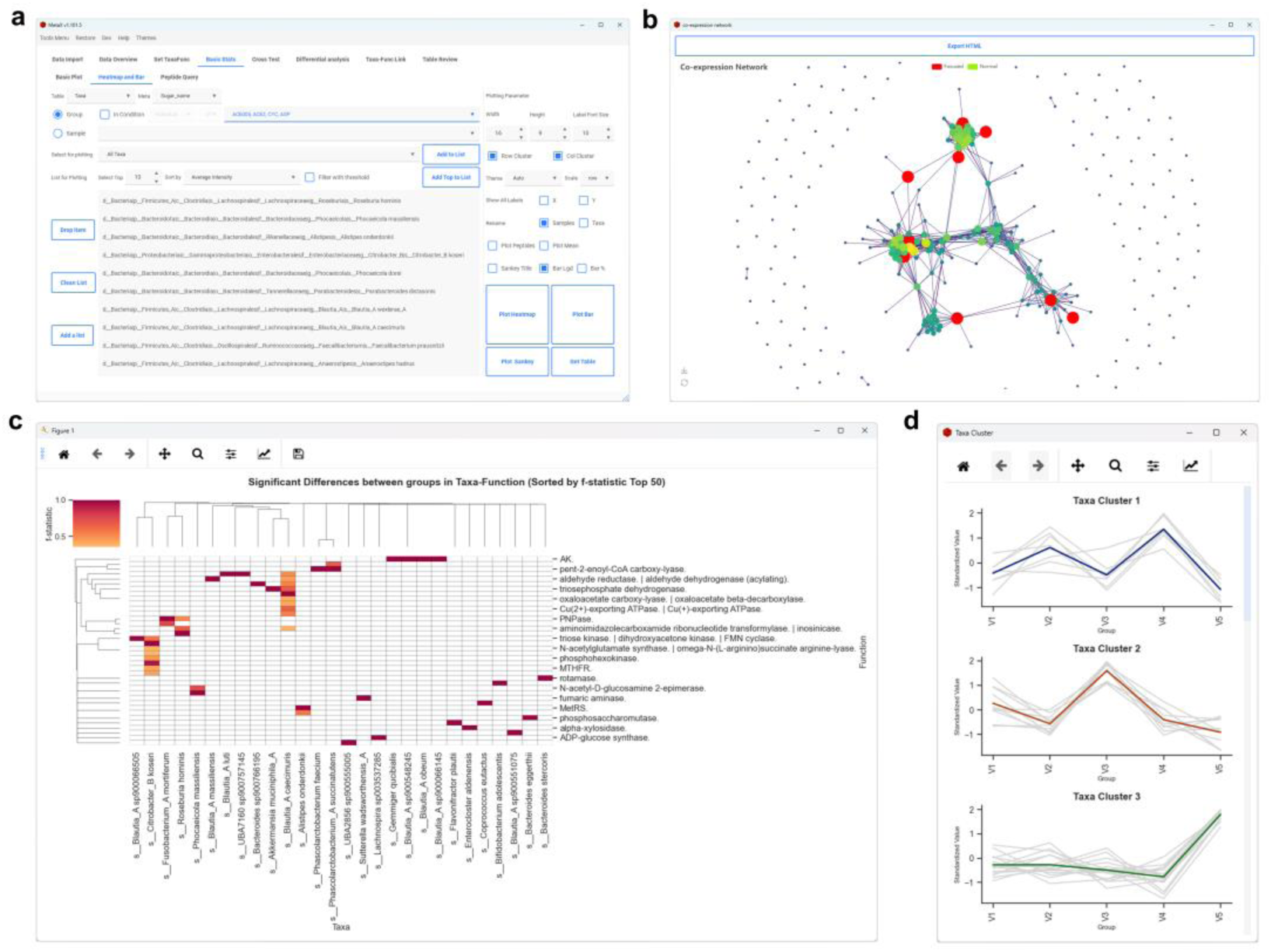
(a) Basic Statistics Interface of MetaX. (b) Co-expression Network Visualization. (c) Heatmap of OTFs Significance. (d) Expression Trend Clusters.

An interesting feature is that for the Tukey Test, MetaX enables a focused examination of all taxa within a selected function or all functions within a chosen taxon. If both a function and a taxon are selected, the software extends the analysis to encompass all peptides within those parameters, with the findings presented in a new window for detailed exploration. This approach facilitates a thorough investigation of the relationships and differences, with options available for visualizing the results through customized plotting. MetaX incorporates Dunnett’s test and Deseq2, which facilitate setting a control group and comparing it against other groups. This functionality is enhanced by the ability to select specific sub-meta groups for more nuanced comparisons under various conditions. Such detailed comparative analysis is crucial for understanding for example treatment effects against control groups within individual subgroups, ultimately leading to an aggregated presentation of results for comprehensive plotting and interpretation. Users have the flexibility to export all result tables and graphs from MetaX. Moreover, all HTML figures are interactive.

For quantitation, MetaX allows users to compute the log2 fold change between two selected groups directly from the Fold Change calculation tab. Upon computation, the results are displayed in a new window, with entries added to a selection list for further analysis. Users have the option to generate Volcano and Sankey plots, adjusting parameters like *p-value*, *p-adjust*, minimum log2 fold change (min-log2FC), and maximum log2 fold change (max-log2FC). Notably, for OTFs table results, the Sankey plot visually connects functions to their respective taxa nodes, facilitating an intuitive understanding of the data’s taxa and function interaction relationships.

MetaX also performs co-expression network analysis. This feature enables the examination of relationships between taxa, functions, peptides or OTFs using either Pearson or Spearman correlation coefficients. Users can define a correlation threshold for plotting the co-expression network. A distinctive feature allows for the marking of certain items as focal points within the network, or for plotting solely those items related to co-expression. This functionality extends to include top-ranked items based on specified criteria, enhancing targeted analysis (illustrated in Fig. 3b).

#### Expression Trends Cluster Analysis

In this analytical module, MetaX empowers users to individually explore expression trends within separate datasets. By selecting specific groups or samples for analysis, users can identify distinct clusters that exhibit similar expression patterns. The resultant clusters are then visualized in a unified figure (Fig. 3d), with the capability to further explore individual clusters via interactive HTML bar or line plots, offering a detailed perspective on expression dynamics.

### Linking the Taxa and Functions

MetaX was tested using the metaproteomics raw files from Sun et al., which utilized LC-MS/MS techniques to explore the gut microbiomes of five individuals exposed to 21 sugar substitute sweeteners. Utilizing MetaLab-MAG software, our study identified 182,370 peptides from the comprehensive dataset comprising 175 samples. Subsequent processing with MetaX software refined the dataset to 174,475 peptides after implementing quality control measures. Of these, 85,775 peptides (49.08% of the dataset) were annotated at the genome level, 65 peptides (0.04%) at the species level, and 40,070 peptides (22.93%) at the genus level, all maintaining an LCA proportion threshold of 100% (Fig. 4a). Through this analytical process, a total of 899 genomes associated with 763 distinct species were annotated, adhering to a minimum criterion of three peptides for each taxon identification (Fig. 4c).

**Figure 4.**
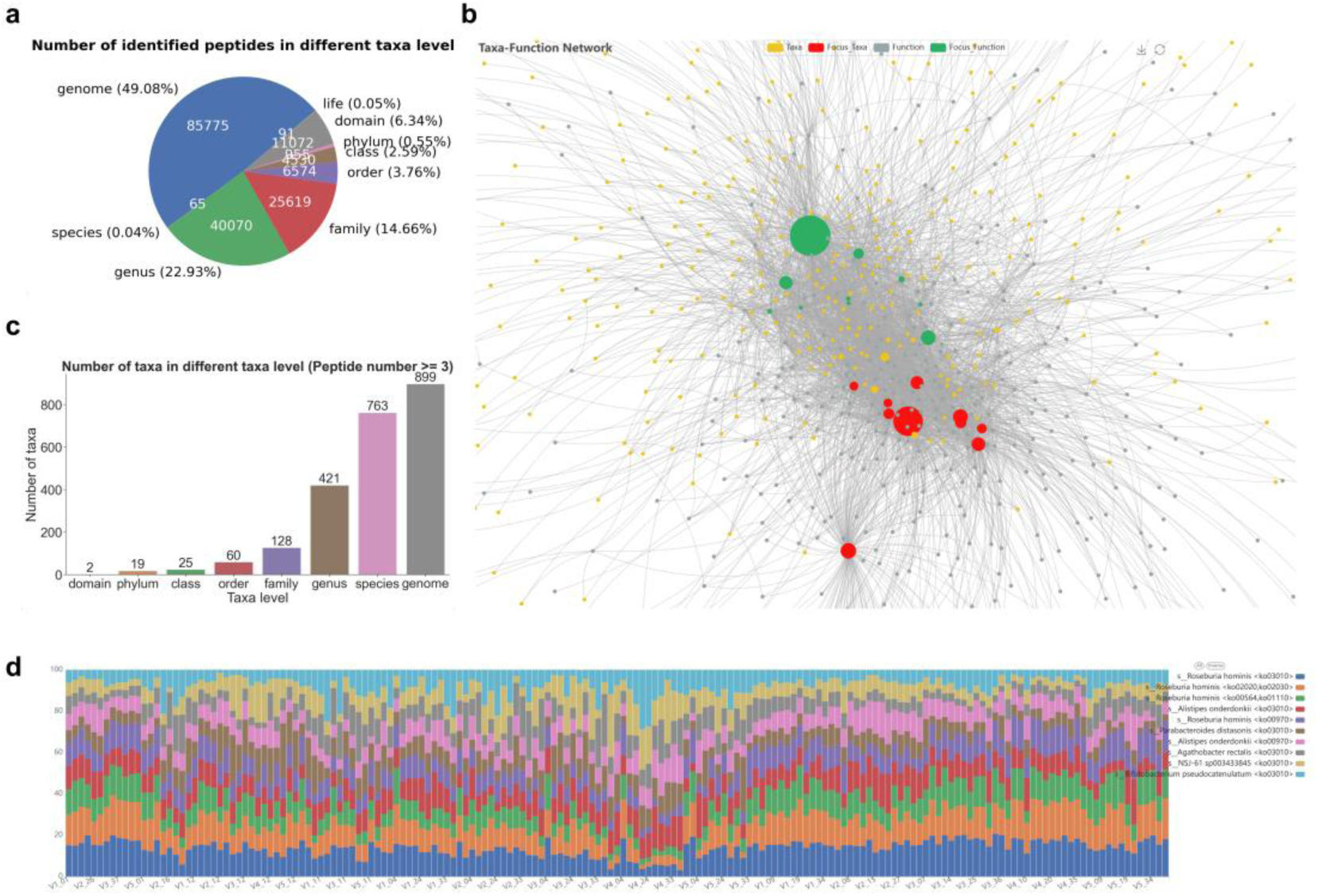
Overview of Test Data. (a) The number of peptide annotations at each taxonomic level. (b) The taxa-functions interaction network. (c) The number of taxa in different levels. (d) Peptide composition in samples within *Roseburia hominis* amino and nucleotide sugar metabolism pathways.

In terms of functions, all peptides were subjected to annotation against multiple databases, including eggNOG (evolutionary genealogy of genes: Non-supervised Orthologous Groups), COG (Clusters of Orthologous Genes), Pfams (Protein families database), GO (Gene Ontology), EC (The Enzyme Commission) and KEGG (Kyoto Encyclopedia of Genes and Genomes), among others. This process revealed significant functional annotation, for instance, 136,566 peptides (78.47% of the annotated group) were found within the COG database with a functional proportion of 100%. Comparable proportions of functional identification were observed across the other databases.

For the examination of OTFs, we utilized the KEGG pathway with a stringent 100% function threshold. The analysis was conducted at the species level, requiring at least three peptides per taxon for inclusion. This approach resulted in the identification of 49,613 peptides, including 511 functions, 832 species, and 4,278 associations between taxa and functions. The resultant network diagram (Fig. 4b) prominently features *Roseburia hominis* with the highest number of function connections, whereas the Ribosome displayed the most extensive taxa links. Within this framework, *R. hominis* is associated with 145 diverse functions, including ABC transporters and aminoacyl-tRNA biosynthesis, with certain functions such as carbapenem biosynthesis being group-specific. Similarly, the Ribosome function is connected to 237 taxa, with a significant peptide count attributed to *R. hominis* and other taxa. Figure 4d displays the top ten intensities of OTFs, with *R. hominis* associated with ko03010 (Ribosome) exhibiting the highest intensity across most samples.

In the study by Sun et al. focused on protein level analysis, distinct functional disparities were observed within the Clostridia class between the carbohydrate (CHO) and noncaloric artificial sweetener (NAS) groups, particularly in COG categories such as C (energy production and conversion), J (translation), O (posttranslational modifications, protein turnover, chaperone functions), E (amino acid metabolism and transport), and G (carbohydrate metabolism and transport) (40). Aligning our investigation with these parameters, we divided the samples into three categories: PBS (Phosphate-buffered saline), CHO, and NAS, selecting the COG categories for functional analysis consistent with the initial study (40). MetaX yielded 57,647 peptides, leading to the identification of 21 COG categories with 59 combinations, encompassing 566 genomes across 497 species, and 2,066 OTFs, with the inclusion criterion set at a minimum of three peptides.

Further, we analyzed the log2 fold changes in OTFs between the CHO and NAS groups, employing a rigorous selection criterion of a p-adjust value less than 0.05 and an absolute log2 fold change greater than 1. Based on these criteria, 40 OTFs were up-regulated and 94 down-regulated in the CHO group in comparison to the NAS group (Fig. 5a-c, Supplementary Table S1). These results are consistent with the COG categories identified in the original study with specific species. Interestingly, our analysis with MetaX provides deeper insights, particularly in mapping specific functions down to the genome level. Notably, the genomes MGYG000002517 and MGYG000000198, corresponding to Roseburia hominis and Enterocloster citroniae respectively, emerged as having the most pronounced changes in OTF between the CHO and NAS groups, offering a more granular understanding of functional dynamics.

**Figure 5.**
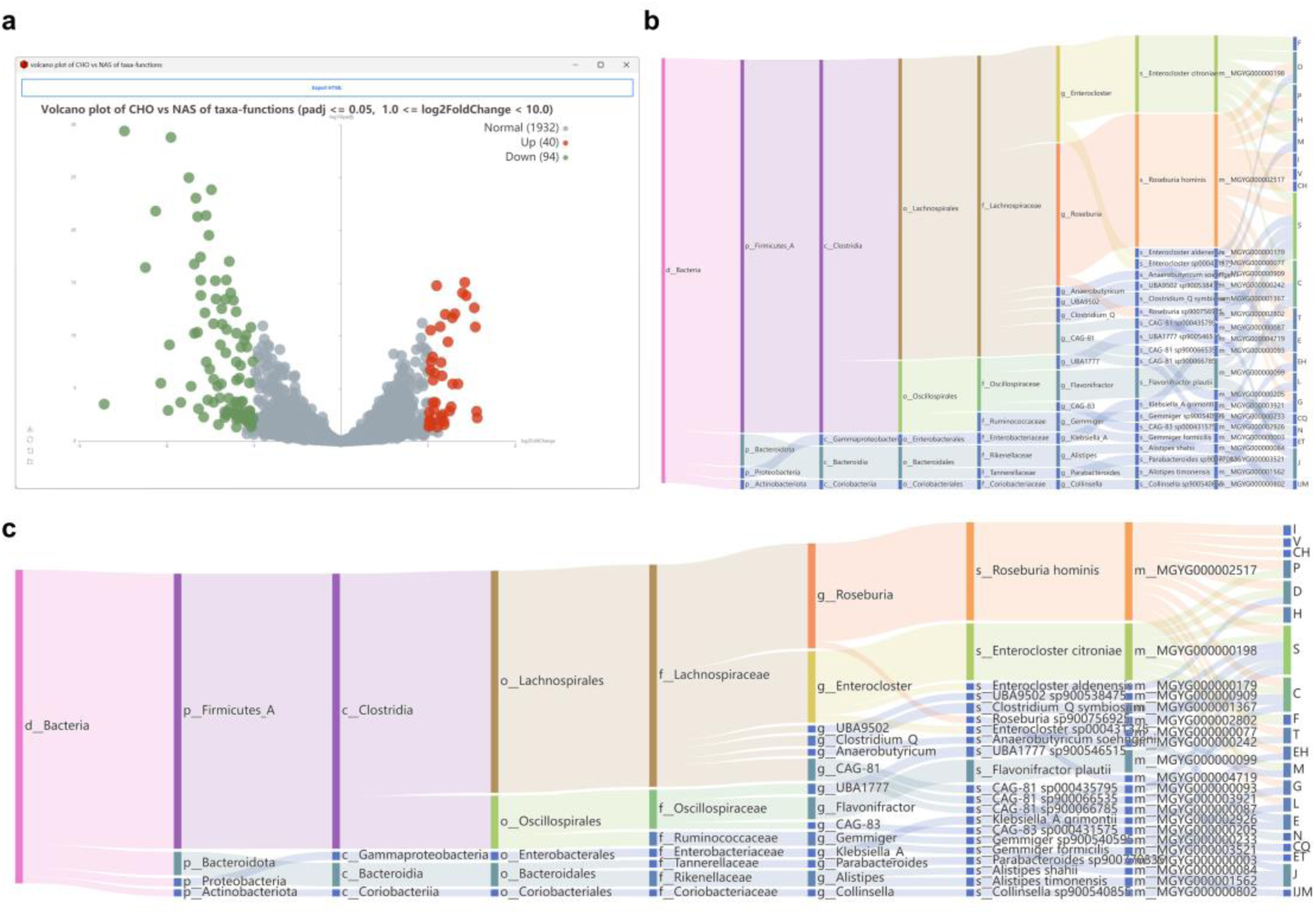
Comparative Analysis of CHO and NAS Groups. (a)Volcano plot of significant difference between CHO and NAS. (b) Sankey diagram detailing taxa-function links for down-regulated OTFs associations between CHO and NAS. (c) Up-regulated OTFs associations between CHO and NAS, with left nodes indicating taxonomic levels and the rightmost nodes denoting functions.

We categorized all sweeteners into distinct groups: sugar alcohols, oligosaccharides, dietary sugars, naturally extracted sweeteners, and synthetic sweeteners. Interestingly, in MetaX, Principal Component Analysis (PCA) revealed distinctive patterns. Specifically, the PCA results for taxa and OTFs indicated that certain extracted sweeteners (Stevioside, Rebaudioside A, Mogroside V) were positioned outside the main cluster. However, this clustering pattern was not observed in the PCA of functions (Fig. 6).

**Figure 6.**
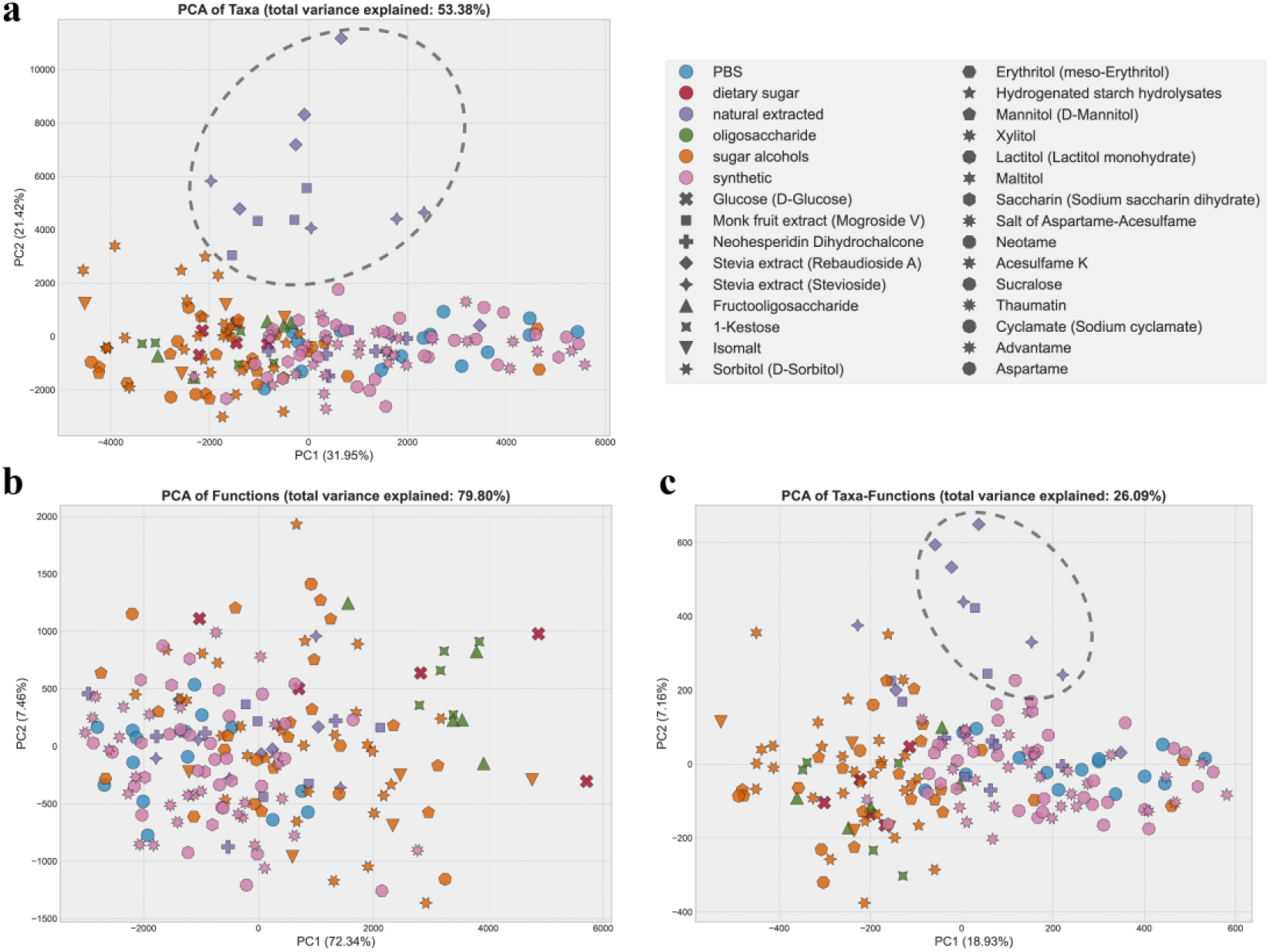
Principal Component Analysis results. (a) PCA results for Taxa. (b) PCA results for functions. (c) PCA results for OTFs.

We then selected the KEGG pathway for analysis, setting the minimum number of linked peptides at three for each taxon, function, and OTF.

First, MetaX was utilized to perform and summarize multiple T-tests, analyzing the OTFs of three sweeteners outside the main cluster in a PCA of taxa and OTFs, compared with PBS. This analysis aimed to identify taxa without significant differences between groups, in contrast to their associated functions which exhibited significant variations, and vice versa. Results with a *P-value* of ≤ 0.01 are detailed in Fig. 7 (Supplementary Table S2). In Fig. 7a, 24 species showed no significant variance between groups, although certain associated functions did. For instance, the abundance of *R. hominis* remained consistent across groups, yet it was linked to 7 functions with significant variations. Conversely, Fig. 7b reveals that 79 functions remained consistent between groups, while the taxa associated with these functions did not. Notably, a significant number of functions associated with *Parabacteroides distasonis* showed substantial differences, underscoring the role of different species in contributing to specific functions across the groups, with *P. distasonis* being particularly pivotal.

**Figure 7.**
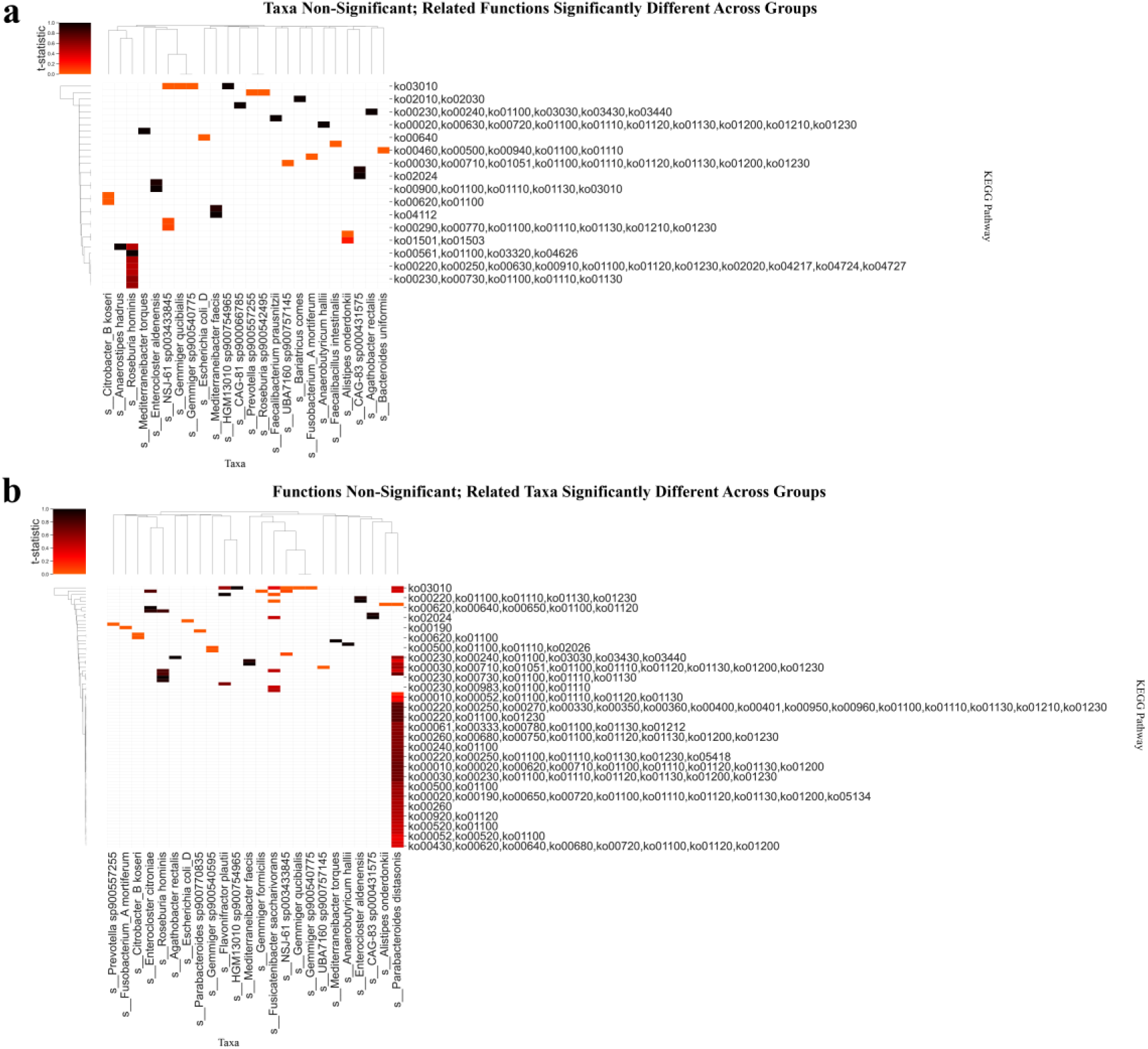
Heatmap of significant OTFs between three sweeteners and PBS: (a) taxa are non-significant but related functions are significant; (b) functions are non-significant but related taxa are significant.

Subsequent analysis using DESeq2 in MetaX compared the taxa, functions, and OTFs of each sweetener to PBS for each individual, respectively. Criteria for significance included a *P-value* of ≤0.05 and |log2FC| ≥1, with consistent trends across individuals (i.e., all individuals exhibited the same significant treatment effect). Interestingly, *P. distasonis* exhibited increased exclusively in Stevioside (STE), Rebaudioside A (REB), and Mogroside V (MFE) across four individuals (V2-V5), aligning with the distinct taxa clustering observed in PCA (Fig. 8a). Further, OTFs analysis identified increased of specific pathways in *P. distasonis* including ko00190 (oxidative phosphorylation), ko00100 (metabolic pathways), ko00230 (purine metabolism), ko00240 (pyrimidine metabolism), ko03020 (RNA polymerase) and others within these groups (Fig. 8c). Yet, no significant functional variations were detected across these conditions, maintaining a consistent trend across all analyzed individuals (Fig. 8b). Additionally, DESeq2 evaluation of CAzy-associated OTFs revealed significant alterations exclusively linked to *P. distasonis* and GH109 across the three sweeteners (Supplementary Table S3).

**Figure 8.**
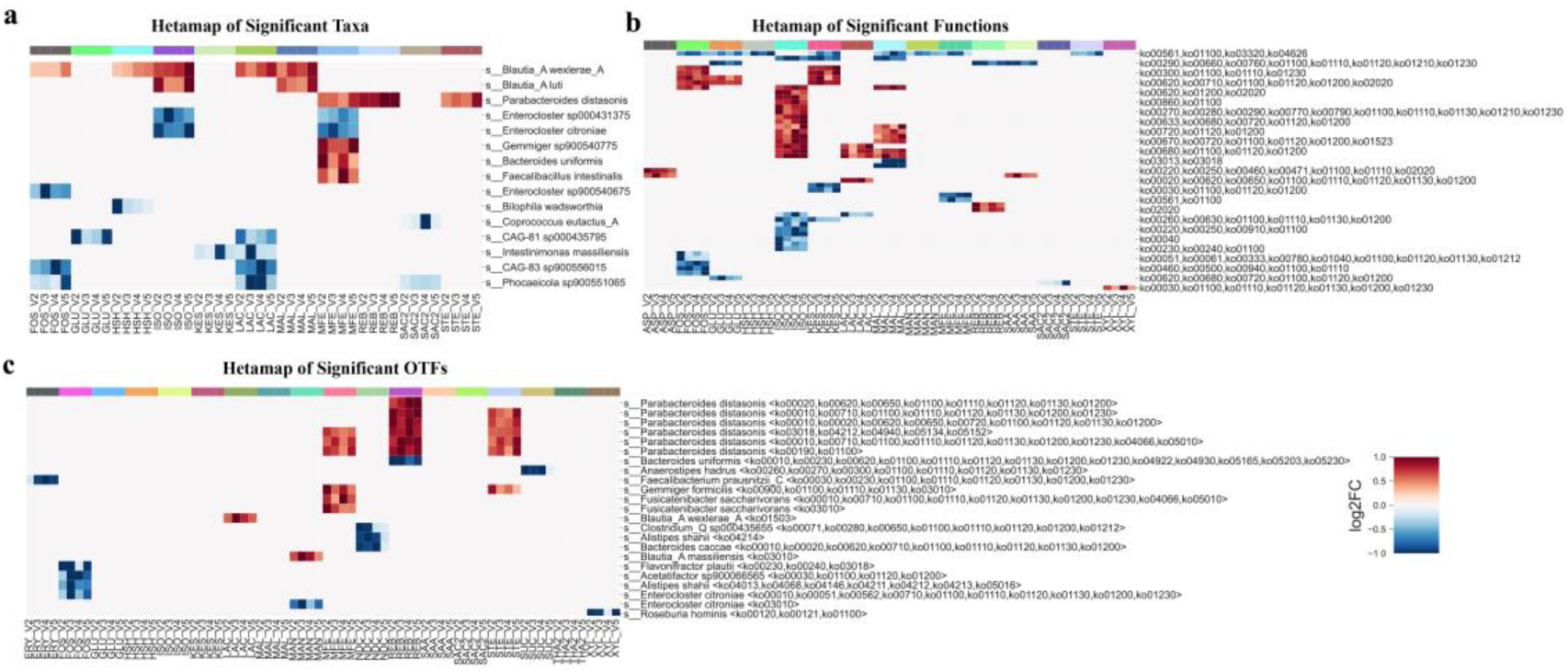
Heatmaps depicting significant changes with consistent trends across four individuals, with columns lacking significant results omitted for clarity (a) Significant taxa. (b) Significant OTFs. (c) Significant functions.

## Discussion

Microbial communities play crucial roles across diverse systems, from human health to environmental nutrient cycling and industrial applications. Their complex interplay of taxonomic structure, functional capabilities, environmental adaptability, and synergistic relationships underpins the necessity for an advanced understanding of these entities (41). Prior investigations into the taxonomic-functional linkage within microbial ecosystems have yielded significant insights and propelled the development of sophisticated computational tools for microbiome analysis (42–44). Current tools, while valuable, are often limited to specific annotations such as GO terms, UniProtKB, or KEGG pathways, and do not provide comprehensive peptide-level metaproteomics annotations and statistical analyses (19–21).

MetaX introduces the OTF framework, a novel conceptual unit that categorizes peptides according to their shared taxonomic and functional characteristics using a proportion-based peptide annotation method. The OTF unit does not restrict taxa to specific functional types. MetaX enables users to create unique OTF pairings by choosing from a variety of taxonomic and functional categories, e.g., Genome-KEGG, Species-GO, Class-EC. It permits the amalgamation of any function with different taxonomic levels, thus expanding the possibility of linking taxa to structural annotations of proteins.

In bottom-up proteomics, peptides are identified by matching MS/MS data to peptide databases generated by *in silico* digestion of proteins. This method relies on matching experimental data with computational predictions(45). Within metaproteomics, the use of metagenome-assembled genomes (MAGs) is common for taxa identification (16). Nevertheless, generating reference-quality, species-level assemblies from metagenomic samples is notoriously challenging (46). To address this, MetaX prioritizes genomes over species as the initial taxonomic level, thereby enabling direct linkage of genomes to functions even in the absence of species-level annotations. Additionally, considering the dynamic nomenclature in microbiology, utilizing genome-level data offers a more accurate reflection of microbial identities(47).

We exemplified the performance of MetaX using a previously published dataset of 175 samples (40). Using MetaX, we not only observed similar results to the protein centric approach previously used but also expanded the analysis to cover a wider range of taxa, achieving insights at the species and genome levels previously unattained. Our findings highlight the versatility of diverse microorganisms in performing similar functions across different environmental conditions, as evidenced in the literature (48–52). MetaX revealed the preferential increase of *P. distasonis* following specific treatments, shedding light on its unique functional contributions within the microbiome. It was through the application of the OTF framework that we were able to discern the specific increase of *P. distasonis* and its associated metabolic functions in response to certain sweeteners: Stevioside, Rebaudioside A, and Mogroside V. This observation not only highlights the intricate interplay between diet and microbial functionality but also emphasizes the critical role of the OTF framework in linking taxonomic data with functional insights. The increase of *P. distasonis* suggests a targeted microbial response, wherein this bacterium proliferates and activates specific metabolic pathways when exposed to these dietary components. The role of *P. distasonis* in the gut microbiota is profound, influencing various health aspects, including antimicrobial resistance and disease modulation (53–59), illuminating its significance in dietary-microbiome interactions and potential implications for human health.

## Conclusion

This study introduces a significant innovation in metaproteomic analysis through the OTF framework and a proportion-based peptide annotation method, linking microbial taxa and functions more effectively. With enhanced visualization tools, we make metaproteomics accessible to a broader audience. Our findings highlight the impact of specific dietary sweeteners on microbial responses, emphasizing the concept of functional redundancy in the gut microbiome and the potential for dietary modulation of gut health.

## Availability and requirements

**Project name:** MetaX

**Project home page:** https://github.com/byemaxx/MetaX

**Operating system(s):** Windows, MacOS and Linux

**Programming language:** Python

**Other requirements:** Python 3.7 or Higher

**Any restrictions to use by non-academics:** Licence needed.

MAGs: metagenome-assembled genomes
UHGP: Unified Human Gastrointestinal Protein
GUI: Graphical User Interface
LC-MS/MS: Liquid Chromatography Coupled with Tandem Mass Spectrometry
PSM: Peptide Spectrum Matches
GO: Gene Ontology
EC: Enzyme Commission number
CAZy: Carbohydrate-Active enZymes
LCA: The Lowest Common Ancestors
CLI: Command-Line Interface
IQR: Interquartile Range
KNN: K-Nearest Neighbors Algorithm
PCA: Principal Component Analysis
ANOVA: Analysis of Variance
eggNOG: Evolutionary genealogy of genes: Non-supervised Orthologous Groups
COG: Clusters of Orthologous Genes
Pfams: Protein families database
EC: The Enzyme Commission
KEGG: Kyoto Encyclopedia of Genes and Genomes
PBS: Phosphate-buffered saline
CHO: Carbohydrate
NAS: Noncaloric artificial sweetener

## Declarations

The authors declare that they have no competing interests.

## Acknowledgements

The authors gratefully acknowledge the assistance of ChatGPT, Grammarly, and DeepL in enhancing the English language presentation of this manuscript. Additionally, certain graphical elements within the overview figures were created with the aid of BioRender.

