## supplementary Document for "MetaX: A peptide centric metaproteomic data analysis platform using Operational Taxa-Functions (OTF)"

### 1. The method description of data preprocessing

**Table 1**. The method of outlier detection

| Name | Description |
|---|---|
| IQR | In a group, if the value is greater than Q3+1.5*IQR or less than Q1-1.5*IQR, the value will be marked as NA. |
| Half-Zero | This rule applies to groups of data. If more than half of the values in a group are 0, while the rest are non-zero, then the non-zero values are marked as NA. Conversely, if less than half of the values are 0, then the zero values are marked as NA. If the group contains an equal number of 0 and non-zero values, all values in the group are marked as NA. |
| Zero-Dominant | This rule applies to groups of data. If more than half of the values in a group are 0, then the non-zero values are marked as NA. |
| Zero-Inflated Poisson | This method is based on the Zero-Inflated Poisson (ZIP) model, which is a type of model that is used when the data contains a lot of zeros, more than what is expected in a standard Poisson model. In this context, the ZIP model is used to detect outliers in the data. The process involves fitting the ZIP model to the data and then predicting the data values. If the predicted value is less than 0.01, then the data point is marked as an outlier (NA) |
| Negative Binomial | This method is based on the Negative Binomial model, which is a type of model used when the variance of the data is greater than the mean. Similar |

| | |
|---|---|
| | to the ZIP method, the Negative Binomial model is fitted to the data and then used to predict the data values. If the predicted value is less than 0.01, then the data point is marked as an outlier (NA). |
| Z-Score | Z-score is a statistical measure that tells how far a data point is from the mean in terms of standard deviations. Outliers are often identified as points with Z-scores greater than 2.5 or less than -2.5. |
| Mahalanobis Distance | Mahalanobis distance measures the distance between a point and a distribution, considering the correlation among variables. Outliers can be identified as points with a Mahalanobis distance that exceeds a certain threshold. |

**Table 2**. The method of handling outlier

| Name | Description |
| --- | --- |
| Original | Outliers will be filled by original value (Remove rows only contain NA and 0 after Outliers Detection). |
| Drop | Discards all rows containing outliers. |
| Mean | Outliers will be imputed by the mean. |
| Median | Outliers will be imputed by the median. |
| KNN | Outliers will be imputed by KNN (K=5). The K-Nearest Neighbors algorithm uses the mean or median of the nearest neighbours to fill in missing values. |
| Regression | Outliers will be imputed by using the IterativeImputer of sklearn with regression method. This method uses round-robin linear regression, modelling each feature with missing values as a function of other features, in turn. |
| Multiple | Outliers will be imputed by using IterativeImputer with multiple imputations method. It uses the IterativeImputer with a specified number (K=5) of the nearest features. |

### 2. Comparing the structures between sweeteners

This study utilized the ChemMine tools to calculate the atom pair and maximum common substructure (MCS) similarities among sweeteners, employing the Tanimoto coefficient as the measure of similarity. Subsequently, multidimensional scaling (MDS) analysis was applied to visualize the relationships between the sweeteners. The MDS plot revealed that three sweeteners, namely Stevioside, Rebaudioside A, and Mogroside V, clustered together (Fig. 1).

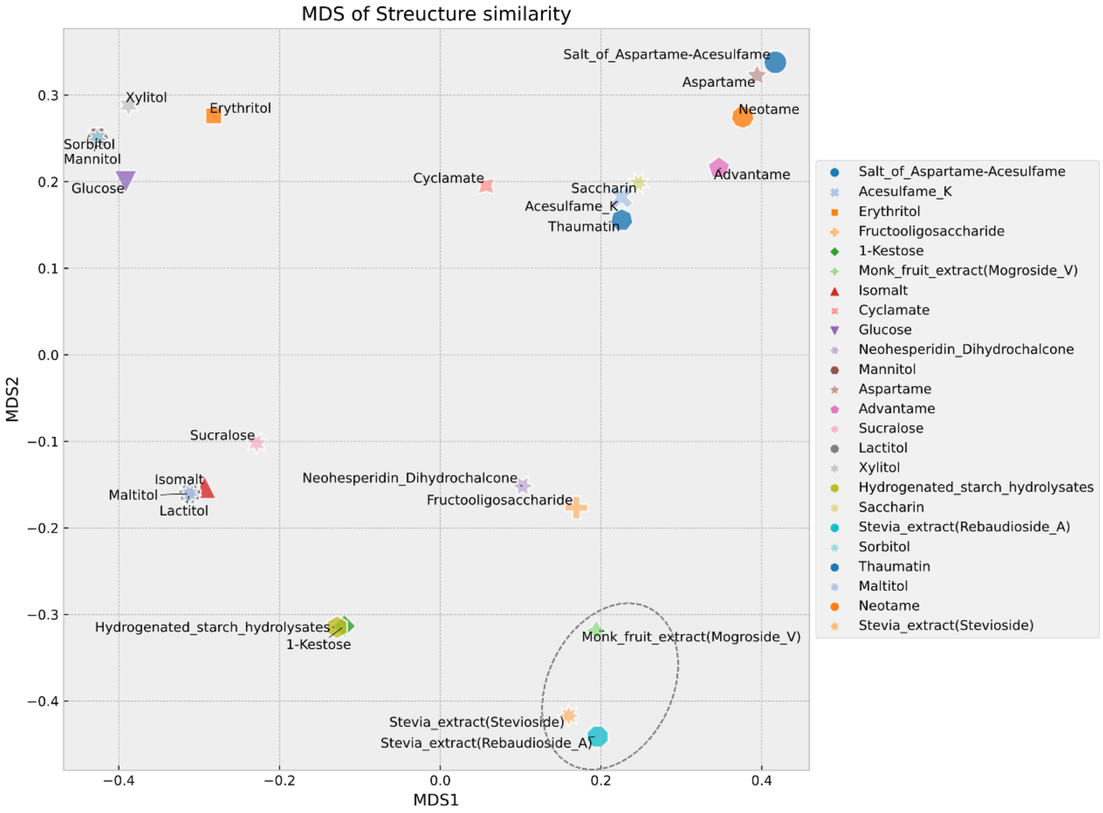


**Figure 1.** Multidimensional Scaling Plot of Structural Similarity Among Sweeteners

Further, physicochemical descriptors for the compounds were computed using the Open Babel software library. The resulting cluster map also grouped these three sweeteners into a single cluster (Fig. 2).

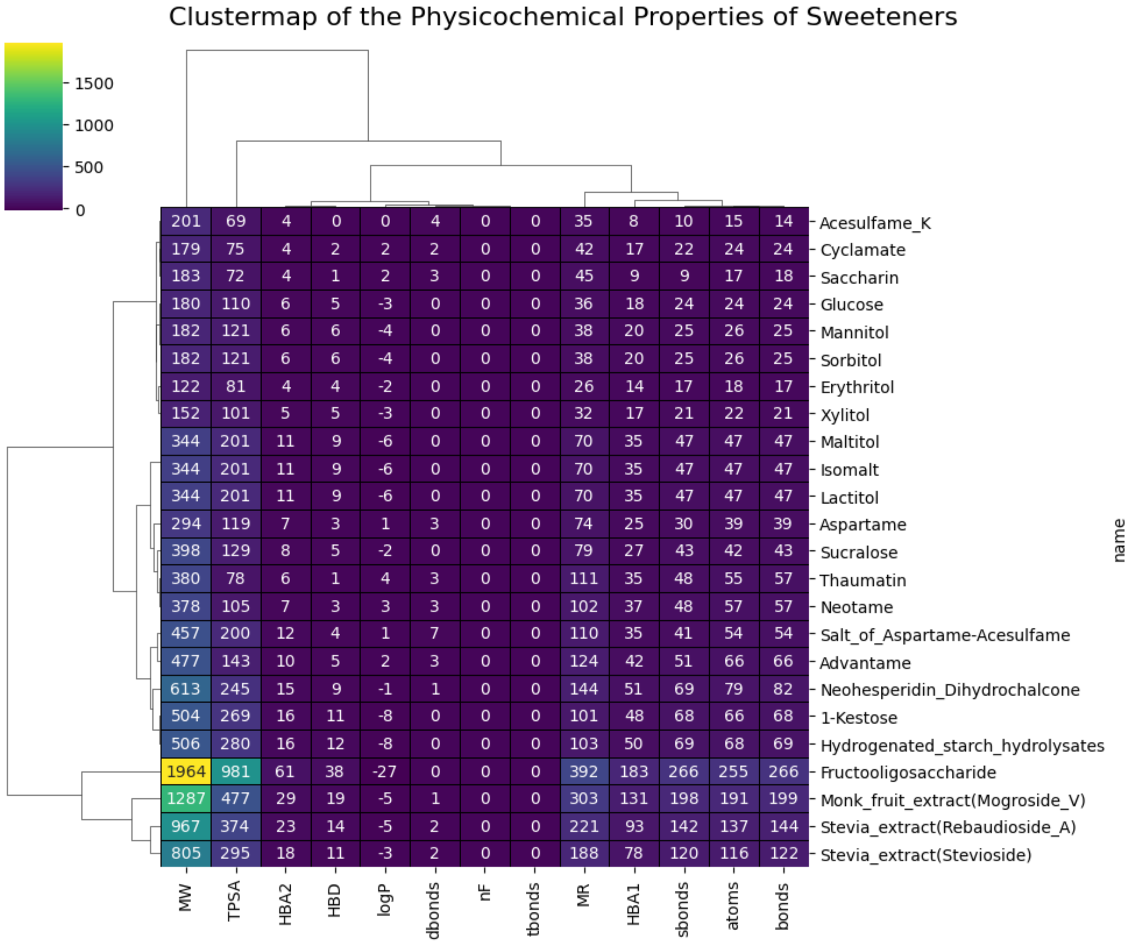


**Figure 2.** Cluster Map of Physicochemical Properties of Sweeteners

A detailed structural comparison of the three sweeteners was conducted by examining atom pairs and the maximum common substructure. This analysis revealed that the sweeteners share structural similarities, including multiple ring structures and several hydroxyl groups. Notably, each molecule contains at least one pentacyclic isoprene unit, which is a long carbon chain with two methyl branches, as part of their side chains (Fig. 3).

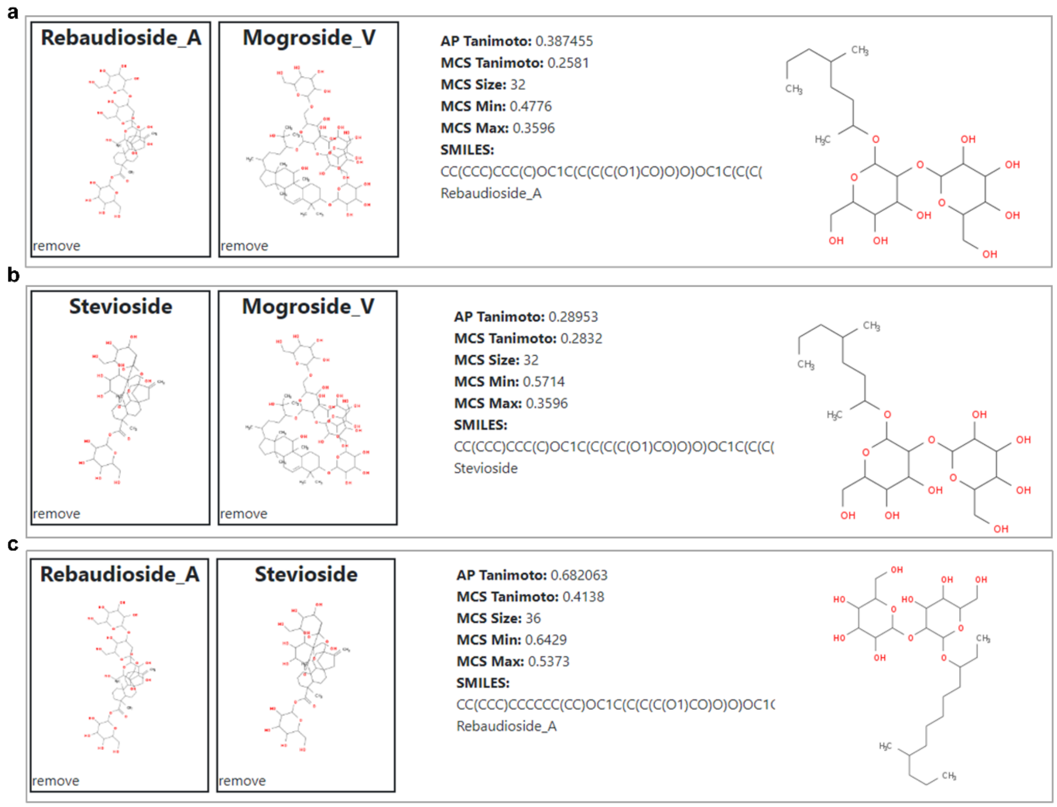


**Figure 3.** Comparative Analysis of Compound Similarities Among Sweeteners. (a) Similarity between Rebaudioside A and Mogroside V. (b) Similarity between Stevioside and Mogroside V. (c) Similarity between Rebaudioside A and Stevioside.
